# Single-Amplicon Sequencing of Coding-Complete Enteroviruses and Rhinoviruses Genomes from Clinical Specimens

**DOI:** 10.64898/2026.10.19.760066

**Authors:** Stephanie Goya, Preston C Longley, Clarice Mauer, Alexander Greninger

## Abstract

Rhinoviruses and enteroviruses are among the most frequently detected respiratory viruses, yet routine diagnostics do not report species or genotype, and standard VP1 genotyping cannot resolve transmission chains or recombination. Recovering sequence beyond VP1 currently requires probe-based enrichment, deep metagenomic sequencing, or several overlapping amplicons. We developed a single-amplicon PCR and sequencing workflow that generates coding-complete genomes from both enteroviruses and rhinoviruses in a single reaction. We evaluated the protocol on 140 rhinovirus-positive respiratory specimens and a panel of 41 isolates spanning 26 enterovirus A-D types, comparing it with hybridization capture sequencing and relating its performance to viral load measured by RT-qPCR and droplet digital RT-PCR. Single-amplicon sequencing recovered 116 high-quality rhinovirus genomes, across 57 rhinovirus genotypes, compared with 113 from hybridization capture. Single-amplicon sequencing was associated with higher fidelity and more uniform sequencing depth compared to hybridization capture. The same primer configuration also recovered two unexpected coxsackieviruses (A6 and A16) from rhinovirus-positive clinical specimens, and the expected polyprotein-coding sequences from all 40 enterovirus isolates tested. Recovery increased with viral abundance, and a visible amplicon band predicted sequencing success. Single-amplicon sequencing reagents cost approximately three-fold less per sample than hybridization capture ($30 versus $85) and required one-third as much time. These findings support single-amplicon sequencing as a simple, cost-effective approach for genomic surveillance of enteroviruses and rhinoviruses from clinical specimens.

**Importance:** Routine respiratory virus tests can detect rhinoviruses and enteroviruses but seldom identify the circulating viral lineage, while VP1 genotyping of rhinoviruses and enteroviruses cannot adequately resolve transmission or recombination. Genome sequencing can provide this information, but existing approaches may require extensive sequencing, probe-based enrichment, or multiple overlapping PCR products. We describe a single-round, single-amplicon method that recovers the polyprotein-coding region across both rhinoviruses and enteroviruses. The method produced highly uniform coverage and reproducible consensus sequences at approximately one-third the price of hybridization capture, and it includes a simple pre-sequencing quality check. This streamlined workflow can help make enterovirus and rhinovirus genomic surveillance more accessible to clinical and public health laboratories.

## Introduction

The genus *Enterovirus* (EV) contains 15 species, including seven human-associated species (formerly *Enterovirus A-D* and *Rhinovirus A-C*, now *Enterovirus alphacoxsackie, E. betacoxsackie, E. coxsackiepol, E. deconjuncti, E. alpharhino, and E. betarhino, E. cerhino,* respectively) that circulate continuously in people and cause illness ranging from common colds to aseptic meningitis, myocarditis, and acute flaccid myelitis (1, 2). Their public health impact reflects both their high burden of infection – enteroviruses routinely comprise the plurality of respiratory virus detections – as well as the emergence, spread, and genetic diversification of individual viral lineages such as poliovirus, EV-D68, and EV-A71 (3–6). Routine diagnostic assays determine whether an enterovirus is present but generally do not identify which enterovirus or rhinovirus lineage is spreading (7–10). Within a single geographic area and time period, detected cases may represent a single transmission chain, repeated introductions of related viruses, or co-circulation of distinct enterovirus species and genotypes (9). Although VP1 genotyping, typically based on an approximately 900 nt region, enables species and genotype assignment, sequence data beyond VP1 are needed to resolve transmission structure, evolution dynamics, and recombinant genomes.

Laboratories pursuing enterovirus genomic characterization generally rely on either untargeted metagenomic sequencing or targeted approaches based on hybridization capture or amplicon sequencing (11, 12). Metagenomic sequencing does not require prior assumptions about which virus or lineage is present and is valuable for viral discovery (13, 14). However, enterovirus genome recovery depends on the proportion of viral reads relative to host and microbial background. At low viral loads, viral reads may represent only a small fraction of the total, requiring higher sequencing depth, which can add cost. Hybridization capture enriches the viral fraction improving recovery of genomes from low viral load specimens. However, recovery is bounded by the reference sequences utilized for probe design: divergent genotypes, genotypes with few or no public reference genomes, and recombinants may enrich poorly. Capture probes, overnight hybridization, and additional library preparation steps also increase the per-sample cost. The third approach is enrichment via targeted reverse transcription polymerase chain reaction (RT-PCR) amplification. Published protocols for near complete genome sequencing from enteroviruses generally use several overlapping amplicons to tile the genome and have recovered diverse genotypes from clinical material (15, 16). However, increasing the number of amplicons also increases the number of primers, reaction complexity, and the risk of uneven amplification across the genome (17).

Single-amplicon recovery of the complete polyprotein-coding region has been described for specific enterovirus species, but no protocol has been adapted specifically for the genetic diversity presented by both enteroviruses and rhinoviruses to date (18, 19). Rhinoviruses (RV) present an important challenge: they are among the most frequently detected respiratory viruses, and their three species comprise 168 genotypes, including 81 RV-A, 32 RV-B, and 55 RV-C, with relatively low nucleotide identity (50-80%) between genotypes across the polyprotein-coding region (20, 21). In one geographically defined population, we observed up to 30 RV-A genotypes co-circulating in a single month; across 8 months of surveillance, we identified an A105/A21 recombinant lineage with a likely breakpoint in the nonstructural protein coding region (6).

In this study, we describe a universal, single-round, single-amplicon RT-PCR covering the enterovirus and rhinovirus coding-complete genome. We first evaluated primer conservation in silico across the genus and assay performance on a panel of cultured enterovirus A-D isolates and then characterized performance in primary specimens using a rhinovirus-positive clinical cohort spanning the diagnostic viral load range. We compared genome recovery to viral load measured by quantitative RT-PCR (RT-qPCR) and droplet digital RT-PCR (ddRT-PCR), assessed reproducibility, and compared workflow costs with metagenomic and hybridization capture sequencing.

## Materials and Methods

### Viral specimens and RNA extraction

Enterovirus species A-D isolates were obtained from remnant culture specimens from UW Virology and from the Centers for Disease Control and Prevention (CDC, courtesy of Dr. Shur-Wern Chern). In total, 41 non-rhinovirus enterovirus specimens were evaluated, including two extracted RNA samples obtained from BEI Resources (ATCC): enterovirus D68 RNA (NR-49138) and enterovirus A71 RNA (NR-4960). The panel comprised enterovirus A isolates (coxsackievirus A2, A6, A8, and A16, and enterovirus A71); enterovirus B isolates (coxsackievirus A9, coxsackievirus B1 to B5, echovirus 4, 6, 7, 9, 11, 16, 18, 25, 30, and 31, and enterovirus B75 and B80); enterovirus C isolates (coxsackievirus A24 and enterovirus C99); and enterovirus D isolates (enterovirus D68). The complete isolate list is provided in Supplementary Table S1.

We selected 140 remnant rhinovirus-positive clinical specimens, collected between November 2023 and November 2024 from patients presenting with respiratory symptoms at UW Medicine (Table 1). Specimens were nasal swabs in viral transport medium and bronchoalveolar lavage samples, originally tested with the AdV/hMPV/RV Assay on the Hologic Panther Fusion platform and stored at −80°C until further processing. The University of Washington Institutional Review Board approved this study with a consent waiver.

**Table 1.** Characteristics of the rhinovirus-positive specimens included in this study.

| Characteristic | Study cohort (n = 140) |
| --- | --- |
| Age, years, median (IQR) | 50 (27-63) |
| Age range, years | 0-85 |
| Male | 89 (63.6%) |
| Female | 51 (36.4%) |
| Nasal swab specimens | 126 (90.0%) |
| Bronchoalveolar lavage (BAL) specimens | 14 (10.0%) |
| Median Hologic Panther Fusion Ct (IQR) | 28.2 (23.6-32.2) |
| Hologic Ct range | 15.8-39.8 |
| Median CDC RT-qPCR Ct (IQR)* | 24.5 (20.6-28.4) |
| Range CDC Ct | 14.6-38.9 |
| Median ddRT-PCR copies/μL RNA (IQR)** | 108.6 (10.5-1,243.9) |
| Range ddRT-PCR copies/μL RNA | 0.23-48,739.8 |
| Samples with RV genotype assigned (%) | 128 (91.4%) |
| Distinct RV genotypes represented | 58 |
| RV-A genotypes | 26 |
| RV-B genotypes | 10 |
| RV-C genotypes | 22 |

\* CDC RT-qPCR Ct was available for 135/140 specimens; the remnant 5 specimens resulted in no detected signal.
\*\* Quantifiable ddPCR results were available for 126/140 specimens; samples below the assay detection/quantification criterion were excluded from calculation of the concentration summary.

The Hologic Panther Fusion assay performs nucleic acid extraction on the instrument as part of its integrated workflow. For all other assays, 140 µL of each clinical or cultured specimen was processed using the QIAamp Viral RNA Mini Kit (Qiagen), and RNA was eluted in 60 µL of the manufacturer’s elution buffer. Culture isolates followed the same procedure, except that 10 µL of clarified viral isolate was diluted to a final volume of 140 µL with phosphate-buffered saline (PBS).

### Rhinovirus/Enterovirus RT-qPCR and droplet digital RT-PCR

Rhinovirus RNA from the clinical cohort was quantified by one-step RT-qPCR using the AgPath-ID One-Step RT-PCR Reagents (Thermo Fisher), according to the manufacturer’s instructions. The assay used the RV|EV-specific primer and probe sequences reported by the CDC: forward 5′-CPA GCC TGC GTG GC-3′, reverse 5′-GAA ACA CGG ACA CCC AAA GTA-3′, and probe 5′-FAM-TCC TCC GGC CCC TGA ATG YGG C-3′ (22, 23). Briefly, 20 µL mastermix contained 12.5 µL 2x RT-PCR buffer, 1µL 25x RT-PCR enzyme mix, 2.5 µL forward primer (10 µM), 2.5 µL reverse primer (10 µM), 0.25 µL probe (10 µM) and 1.25 µL nuclease-free water, and was mixed with 5 µL of extracted RNA.

Thermocycling consisted of 10 min at 45°C for cDNA synthesis, followed by 10 min at 95°C and 40 cycles of 15 s at 95°C and 45 s at 60°C, with fluorescence measured at the end of the 60°C step. This assay is referred to hereafter as CDC RT-qPCR.

The same primer and probe set was used for one-step ddRT-PCR with the One-Step RT-ddPCR Advanced Kit for Probes (Bio-Rad). Each reaction contained 5 µL of Supermix, 2 µL of reverse transcriptase, 1 µL of 300mM DTT, 0.9 µL of 10 µM primer forward, 0.9 µL of 10 µM primer reverse, 0.5 µL of 10 µM probe and 9.7 µL of nuclease-free water and 2 µL of extracted RNA, and droplets were generated with the QX 200 AutoDG System (BioRad).

Thermocycling consisted of a reverse transcription incubation of one hour at 50 °C, PCR enzyme activation of 10 min at 95 °C, 40 cycles of 30 s at 95 °C and 1 min at 60 °C with a ramp of 1.5 °C/s, a final enzyme deactivation of 10 min at 98 °C and hold at 4 °C until droplet reading. Amplification products were analyzed on a Bio-Rad QX600 Droplet Digital PCR System. Positive and negative droplets were classified using QuantaSoft Analysis v1.7.4.0917, and target concentrations were calculated using Poisson distribution-based analysis implemented in the Bio-Rad software. Results are reported as copies/µL of extracted RNA or copies/µL per reaction.

### Single-amplicon amplification and sequencing of the enterovirus polyprotein-coding region

The genome amplification strategy was adapted from an approach previously used for norovirus genogroup II (24, 25). Complementary DNA (cDNA) was synthesized from viral RNA in half-volume reactions with the qScript Ultra Flex Kit (QuantaBio) and the T_30_TAG primer (T30TAG: 5′-GACTAGTTCTAGATCGCGAGCGGCCGCCCTTTTTTTTTTTTTTTTTTTTTTTTTTTTTT-3′) (Figure 1A). Briefly, 1 µL of T_30_TAG primer was mixed with 1 µL of 10X GSP enhancer, 3.5 µL of nuclease-free water and 2.5 µL of extracted RNA. The mix was incubated for 5 min at 65 °C and then 2 min at 4 °C, after which 2 µL of 5x qScript Ultra Reaction Mix was added. Reverse transcription was performed at 55 °C for 10 min, followed by 85 °C for 5 min and hold at 4 °C before PCR.

**Figure 1.**
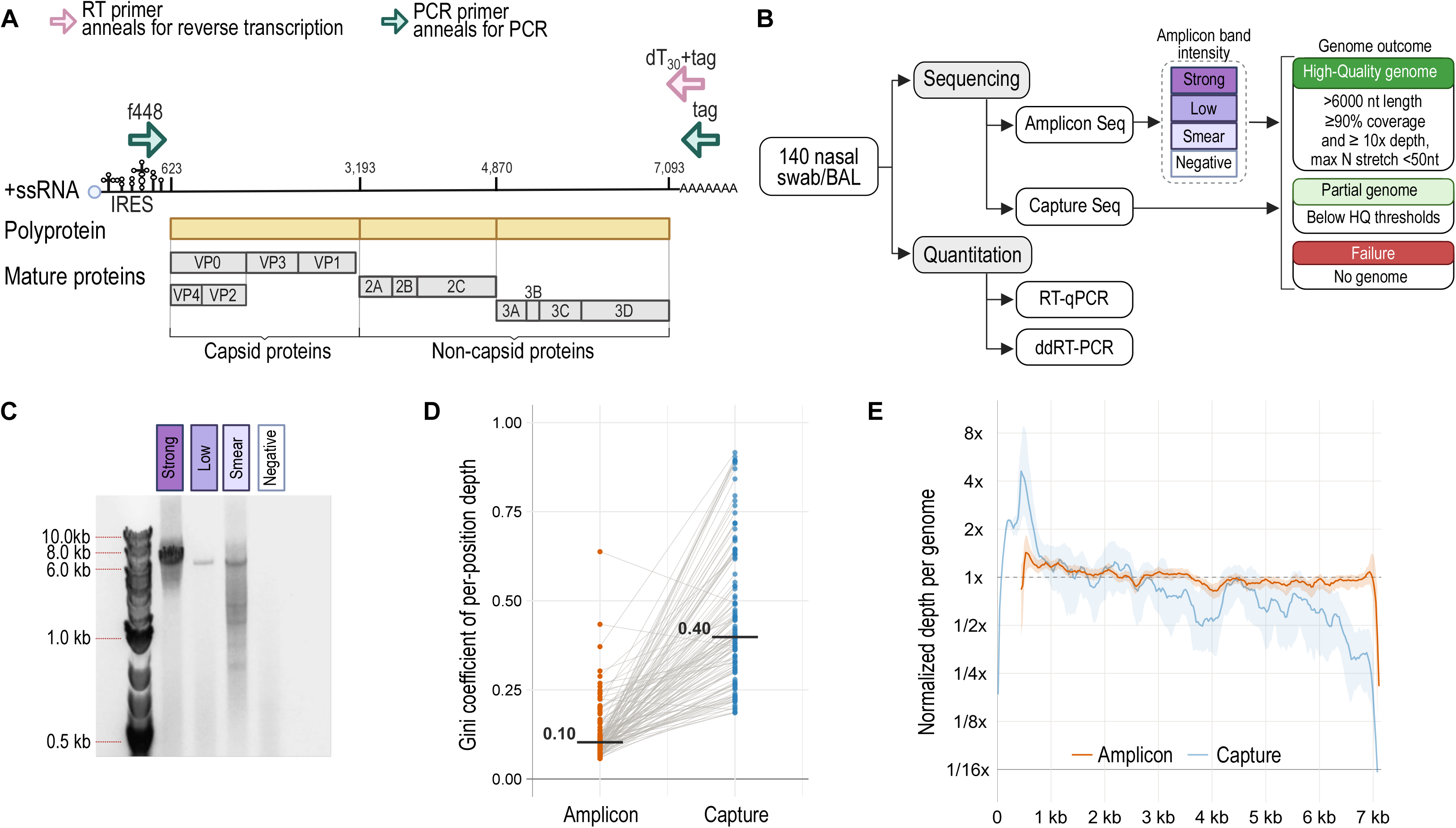
Single-amplicon rhinovirus sequencing strategy, study workflow, and genome-wide depth uniformity. (A) Schematic representation of the positive-sense rhinovirus genome and single-amplicon strategy. Reverse transcription was performed using an oligo(dT)_30_ primer containing a 5′ tag, followed by PCR using the rhinovirus-specific forward primer f448 and a tag-specific reverse primer. Approximate genome coordinates and mature viral proteins are indicated. (B) Study workflow for 140 nasal swab or bronchoalveolar lavage (BAL) specimens analyzed by single-amplicon sequencing, hybridization capture, RT-qPCR, and ddRT-PCR. Amplicon products were categorized as strong, low, smear, or negative based on PCR product electrophoresis results. Recovered sequences were classified as high-quality genomes when they were ≥6,000 nt, had ≥90% of genomic positions covered at ≥10× depth, and contained no N stretch ≥50 nt; other recovered sequences were classified as partial genomes, and samples without a recovered genome were classified as failures. (C) Representative agarose gel illustrating the four amplicon-band categories. (D) Paired comparison of the Gini coefficient of positional depth for 121 sample-genotype pairs recovered by both methods. Lower Gini coefficients indicate more uniform genome-wide coverage. Horizontal bars indicate median values, and lines connect paired genomes. (E) Genome-wide sequencing depth for single-amplicon and hybridization capture. Depth at each position was normalized to the mean depth of the corresponding genome, and genomes were projected onto a standardized 7-kb coordinate system. Lines indicate the median normalized depth and shaded regions the interquartile range. The horizontal dashed line denotes the genome-specific mean depth.

The enterovirus polyprotein-coding region was amplified using a forward primer targeting a conserved region in the 5′ untranslated region of enteroviruses and rhinoviruses (f448: 5′-TCCTCCGGYCCCTGAATGYGGCTAA-3′), and a reverse primer targeting the sequence introduced during reverse transcription (tag: 5′-GACTAGTTCTAGATCGCGAGCGGCCGCCC-3′). PCR was performed in half-volume reactions with repliQa HiFi ToughMix (QuantaBio). The mastermix contained 12.5 µL of repliQa HiFi ToughMix (2X), 0.5 µL of DMSO, 1 µL of forward primer (10 µM), 1 µL of reverse primer (10 µM), 7.5 µL of nuclease-free water, and 2.5 µL of cDNA product was added. Thermocycling consisted of an initial denaturation at 98°C for 30 s, followed by 35 cycles of 98°C for 10 s and 68°C for 40 s. PCR products were visualized by agarose gel electrophoresis with ethidium bromide using 2 µL of the amplification reaction. The remaining 23 µL of PCR product was purified with AMPure XP beads at a 0.45x bead-to-sample ratio and eluted in 30 µL of nuclease-free water.

Purified amplicon concentrations were measured using the Qubit 1x dsDNA High-Sensitivity Assay kit (Invitrogen). Illumina libraries were prepared from up to 100 ng of purified amplicon DNA using half reactions of the Illumina DNA Prep (M) Tagmentation kit (Illumina). Library concentrations were measured by Qubit, and libraries without a quantifiable readout were not included in the sequencing pool. The final pool was checked using the D1000 DNA ScreenTape assay on a TapeStation instrument (Agilent). Libraries were sequenced using either 2×150 bp reads on a NextSeq 2000 instrument or 1×100 bp reads on a NovaSeq6000 instrument (Illumina). No-template controls were included throughout and remained negative.

### Rhinovirus hybridization capture sequencing

Hybridization capture libraries were prepared using 3.4 µL of extracted RNA using the QIAseq xHYB Microbial Hyb & Lib Kit (Qiagen), with each step prior to probe hybridization performed at a two-thirds reaction volume. Viral enrichment was performed with a custom QIAseq xHYB Custom Microbial Panel designed to target all rhinovirus genotypes with complete genome sequences available in the NCBI GenBank up to June 03, 2026. Hybrid capture reactions were performed in pools of up to eight libraries, with a maximum library input of 1500 ng per pool. Libraries were pooled according to the CDC RT-qPCR Ct value of the corresponding re-extracted specimen, with each pool spanning no more than 2 Ct units, to limit differences in estimated viral nucleic acid input among libraries. Library quantification, exclusion of libraries without a quantifiable readout, the final pool check and sequencing (2×150 bp reads on a NextSeq 2000 instrument or 1×100 bp reads on a NovaSeq6000 instrument; Illumina) were performed as for single-amplicon libraries.

### Rhinovirus and enterovirus genome assembly and analysis

Viral consensus sequences were generated with the picorna-strm pipeline (https://github.com/greninger-lab/picorna-strm), which outputs quality-filtered reads, mapping results, coverage metrics, and consensus sequences. Sequencing adapters were trimmed and reads quality-filtered with fastp v0.23.2. Reads were retained if they were longer than 50 nucleotides after trimming and had an average Phred quality score of at least Q20.

Quality-filtered reads were first mapped with BWA-MEM v0.7.19 against the complete picornavirus reference database consisting of complete human picornavirus genomes representing all available species, types, and genotypes in ICTV as of August 2026. Reference genomes with a median genome coverage of at least 30% and an average depth of at least 3x were selected for independent follow-up mapping. Reads were then mapped separately to each selected reference genome, and an intermediate consensus sequence, referred to as ‘Consensus1’, was generated for each mapping in which the reference was covered > 60% with an average depth >5x. ‘Consensus1’ was used as the reference for a final mapping step, and a final consensus sequence was generated for that alignment with over 50% genome coverage and a minimum average depth of 5x. All consensus sequences were generated with iVar v1.4.4. For libraries generated with the single-amplicon protocol, the pipeline was run in amplicon mode, in which the forward primer sequence at the 5′ end of the amplicon was identified and trimmed from the resulting consensus sequence.

The pipeline assigned rhinovirus genotypes based on VP1 nucleotide identity (87% in RV-A and RV-C, 88% in RV-B) against the same August 2026 picornavirus database (26). As a post-assembly quality control, the pipeline compared all consensus genomes within a sequencing run in an all-versus-all manner and flagged pairs from different specimens that share more than 99.9% nucleotide identity, excluding ambiguous positions and gaps, as possible cross-contamination; flagged genomes were manually reviewed and excluded before analysis. Consensus genomes were annotated and quality-verified using VADR (https://github.com/greninger-lab/vadr-vscan-aglab). For sequencing performance analysis, we defined a high-quality genome as ≥90% of nucleotide positions covered at ≥10x depth, total length ≥6,000 nt and the N stretches lower than 50 nt.

Intrahost variability was evaluated only at canonical A, C, G or T consensus discordances. A position was considered evaluable when depth was ≥20x in each compared dataset, and intrahost variation was supported when both consensus alleles were observed and the minor-allele frequency was ≥5% in at least one dataset.

Reproducibility was evaluated within sample-genotype groups rather than treating individual run pairs as independent observations. All consensus genomes from the same sample-genotype group were aligned using MAFFT v7.526. For groups replicated by both sequencing methods, pairwise concordance was first summarized within each sample-genotype group for within-amplicon, within-capture and between-method comparisons, and group-level median concordances were compared across these three comparison types using a Friedman test followed by paired Wilcoxon tests.

### In silico evaluation of forward primer conservation

Forward primer (f448) conservation was evaluated using all nucleotide sequences assigned to genus Enterovirus (NCBI taxid:12059) available through NCBI Virus as of September 29, 2026. Of 140,030 records, 31,128 contained either an explicitly annotated 5′UTR or an unambiguous interval preceding the annotated start of the polyprotein coding sequence. These were aligned with MAFFT v7.526 (--addfragments) against the curated human picornavirus genomic database used by our assembly pipeline (331 complete RV A-C and EV A-D genomes) to locate the 25 homologous primer-binding sites, using an exact-match reference species. Conservation was scored only for sequences with unambiguous coverage of all 25 positions and at least 10 nucleotides in each adjacent 25 nt flank (total sequences: 20,972). The primers of Isaacs et al. and Chien et al. (18, 19) were taken from the original publications and evaluated with the same procedure. For the six nucleotides by which the primer of Chien et al. extends 3′ of f448 primer, conservation was scored as the proportion of sequences carrying a substitution or indel.

### Phylogenetic analyses

All complete human-derived rhinovirus genomes longer than 6,000 nt were downloaded from NCBI Virus on August 27, 2026, using taxonomic identifiers 147711 for RV-A, 147712 for RV-B, and 463676 for RV-C. These reference genomes and the genomes generated in this study were aligned using MAFFT v7.490, and alignments were inspected using AliView v1.28. Maximum-likelihood phylogenetic trees were inferred using IQ-TREE version 2.1, with branch support assessed by ultrafast bootstrap approximation implemented in UFBoot2 with 1,000 replicates. Monophyletic clusters were considered statistically supported when UFBoot2 support was at least 90%. Phylogenetic trees were visualized using FigTree version 1.4.4. Recombination in RV specimens was assessed by comparing the phylogenetic associations inferred from the VP1 and 3D genetic regions. Phylogenetic trees are available in the GitHub repository (https://github.com/greninger-lab/RV_NGS_methods/).

### Statistical analysis

Statistical analyses and figure generation were performed in R version 4.2.2 within RStudio version 2024.04.02; R Markdown files and data frames are available in the study GitHub repository (https://github.com/greninger-lab/RV_NGS_methods/). Continuous variables were compared across groups by Kruskal-Wallis test followed by Dunn post hoc test, or by Wilcoxon signed-rank test when paired. Categorical associations were tested by Fisher exact test, and agreement between quantitative assays by Pearson correlation and Bland-Altman analysis.

Genome recovery was modelled by univariable logistic regression. Model discrimination was quantified by the empirical area under the receiver operating characteristic curve (AUC). Profile-likelihood 95% confidence intervals were calculated for odds ratios. Confidence intervals for probability thresholds were approximated using the delta method and, for models using log10 RNA concentration, back-transformed to the original scale.

Effect sizes were reported as epsilon-squared for Kruskal-Wallis tests, Cramér V for Fisher’s exact test, and the paired Hodges-Lehmann location shift for paired Wilcoxon signed-rank tests. All tests were two-sided, p-values below 0.05 were considered significant, and p-values from multiple comparisons were adjusted by the Benjamini-Hochberg method. Kruskal-Wallis, Wilcoxon, Friedman, Fisher exact, Pearson correlation and logistic regression were implemented with the stats package of base R; the Dunn post hoc test, Cramér V and the Gini coefficient with DescTools v0.99.54. Figures were produced with ggplot2 v3.5.2.

### Use of AI-assisted tools

OpenAI Codex (OpenAI; GPT-5, accessed August 2026) was used to assist in writing code in R and Python for data processing and statistical analysis. All code was reviewed, tested, and debugged by the authors, who defined the analytical approach, executed all analyses on the original data, and verified the outputs. No generative AI tool was used to generate, alter, or interpret data. The authors take full responsibility for the accuracy and integrity of the work.

### Data availability

Raw FASTQ reads generated in this study have been deposited in the NCBI Sequence Read Archive under BioProject accession number PRJNA1521468, and consensus genome sequences in NCBI GenBank. Sample-level accession numbers and associated metadata are provided in Supplementary Table S1.

## Results

### In silico analysis revealed near-universal conservation of the f448 binding site across human enteroviruses and rhinoviruses

Single-amplicon genome sequencing uses a virus-specific forward primer that anneals near the 5′ end of the genome and a reverse primer targeting the poly(A) tail (Figure 1A). The amplicon spans the downstream portion of the 5′ untranslated region (UTR), the polyprotein coding region, and the complete 3′ UTR. The forward primer, f448, derives from a probe previously described in an RT-qPCR assay for rhinovirus detection (23), extended at the 3′ end and including one additional degenerate position. It targets the internal ribosome entry site (IRES) within the 5′UTR, a region expected to be conserved across the genus Enterovirus.

Among 20,972 sequences with complete and unambiguous coverage of the f448 binding region, the primer was an exact match in 20,536 (97.9%), across human rhinoviruses and polio and non-polio enteroviruses (Supplementary Table S2). Mismatches were rare, ranging from 0.58% in EV-D to 3.49% in EV-C, and almost always consisted of a single nucleotide: 385 sequences (1.8%) carried one substitution, 28 (0.1%) two, and 23 (0.1%) three to eight.

The most frequent change was C>A at primer position 22 (22:C>A), found in 138 sequences (25.7% of all substitutions) of EV A-C, RV-A and RV-C. It affected 2.51% of EV-C sequences versus 0.06-0.90% in other species, and was present in most sequences of three infrequently detected respiratory EV-C types: 25/26 (96.2%) EV-C104, 28/37 (75.7%) EV-C105 and 16/16 (100%) EV-C109 (27). Whether a substitution at this position affects amplification efficiency of these enterovirus types cannot be inferred from sequence data alone. Overall, the near-universal conservation of this locus supports f448 as a broadly applicable priming site across human enteroviruses and rhinoviruses.

### Single-amplicon sequencing recovered polyprotein-coding sequences across human enterovirus species A-D

To test f448 on this diversity, we applied our single-amplicon protocol to a panel of 41 enterovirus isolates from human enterovirus species A through D, including coxsackieviruses, echoviruses, EV-A71, EV-C99, and EV-D68 (Supplementary Table S1). The expected genome sequence was recovered from every isolate, and all 32 isolates previously sequenced by shotgun metagenomics yielded identical consensus genomes (Supplementary Table S1). All 41 genomes met the high-quality criteria defined in the Methods, with a median of 97.3% of positions covered at ≥10x (IQR 94.9-100%, minimum 94.0%), and consensus lengths of 6,872-7,343 nt without N stretches (Supplementary Table S1; Supplementary Figure S1).

### Single-amplicon sequencing recovered high-quality rhinovirus genomes from clinical samples

We next evaluated single-amplicon sequencing in 140 remnant RV-positive respiratory specimens previously tested with the AdV/hMPV/RV assay on the Hologic Panther Fusion (126 nasal swabs and 14 bronchoalveolar lavage specimens; Figure 1B). We graded the degree of amplification based on the quality of amplification band after electrophoresis in agarose gel: strong (n = 78), low (10), smear (36), and negative (16) (Figure 1B/C). Single-amplicon sequencing produced genomic data for 118 specimens (84%), including coxsackievirus genomes in two specimens (CV-A6 and CV-A16). Six specimens had co-infections between two rhinovirus species or genotypes and one between rhinovirus and coxsackievirus, giving a total of 124 genomes (Supplementary Table S1). Hybridization capture produced genomes for 127 (91%) specimens, detected the same 5 rhinovirus co-infections (but not the coxsackievirus co-infection) and gave 132 genomes. Using the high-quality definition in the Methods, single-amplicon sequencing recovered 116 high-quality genomes and 6 partial genomes, while hybridization capture recovered 113 high-quality genomes and 19 partial genomes (Supplementary Table S3). High-quality genomes were obtained from libraries with as few as 39,383 quality-filtered reads for single-amplicon and 27,774 for hybridization capture. Capture probes were rhinovirus-specific and did not recover the coxsackieviruses, so the method comparison is limited to rhinovirus.

Single-amplicon sequencing produced significantly more uniform coverage than hybridization capture across the 122 single-amplicon and 132 capture rhinovirus genomes (median Gini coefficient 0.10 [IQR 0.08-0.12] vs 0.40 [IQR 0.30-0.56]; p-value = 2.7 × 10⁻²¹, paired Wilcoxon signed-rank test; Hodges-Lehmann median difference 0.30, 95% CI 0.26-0.35; Figure 1D). The difference in coverage held across all sequenced RV-A, RV-B and RV-C genotypes (Figure 1E, Supplementary Figure S2).

### Single-amplicon sequencing success improves at higher viral abundance, and PCR product visualization identifies the specimens worth sequencing

Viral abundance using three methods, including the diagnostic Panther Fusion Ct value, a CDC RT-qPCR Ct value on re-extracted specimens, and ddRT-PCR copy number (Supplementary Table S3). The methods were concordant (Pearson r = 0.879 between Ct values; Supplementary Figure S3), but the Ct values were not interchangeable: CDC Ct values averaged 2.55 cycles lower than the Panther Fusion (SD 2.66), whereas the 22% lower RNA equivalent used per reaction predicted a Ct 0.36 higher (Supplementary Table S4). The offset was consistent across RV species (mean Panther Fusion minus CDC Ct difference +2.38, +2.96, and +2.91 for RV-A, RV-B, and RV-C; Kruskal-Wallis p-value = 0.15).

Amplicon visualization and DNA quantification provided a helpful screen on the expected yield of sequencing. All 16 specimens without a visible product failed to provide a quantifiable library, while every library that passed quantification yielded a genome. Libraries passed quantification in 78/78 specimens with a strong band, 10/10 with a low-intensity band, and 30/36 with a smear (Fisher exact p = 9.4 × 10⁻⁴, Cramér V = 0.35). Band intensity tracked viral load only coarsely (Kruskal-Wallis P = 7.7 × 10⁻⁹ and 2.5 × 10⁻⁵; Figure 2A, 2B), and low– and smear-band specimens were indistinguishable based on CDC Ct (Dunn adjusted p-value = 0.92) yet yielded genomic data in 100% and 83.3% of specimens, indicating post-amplification gel appearance offered information that Ct values did not.

**Figure 2.**
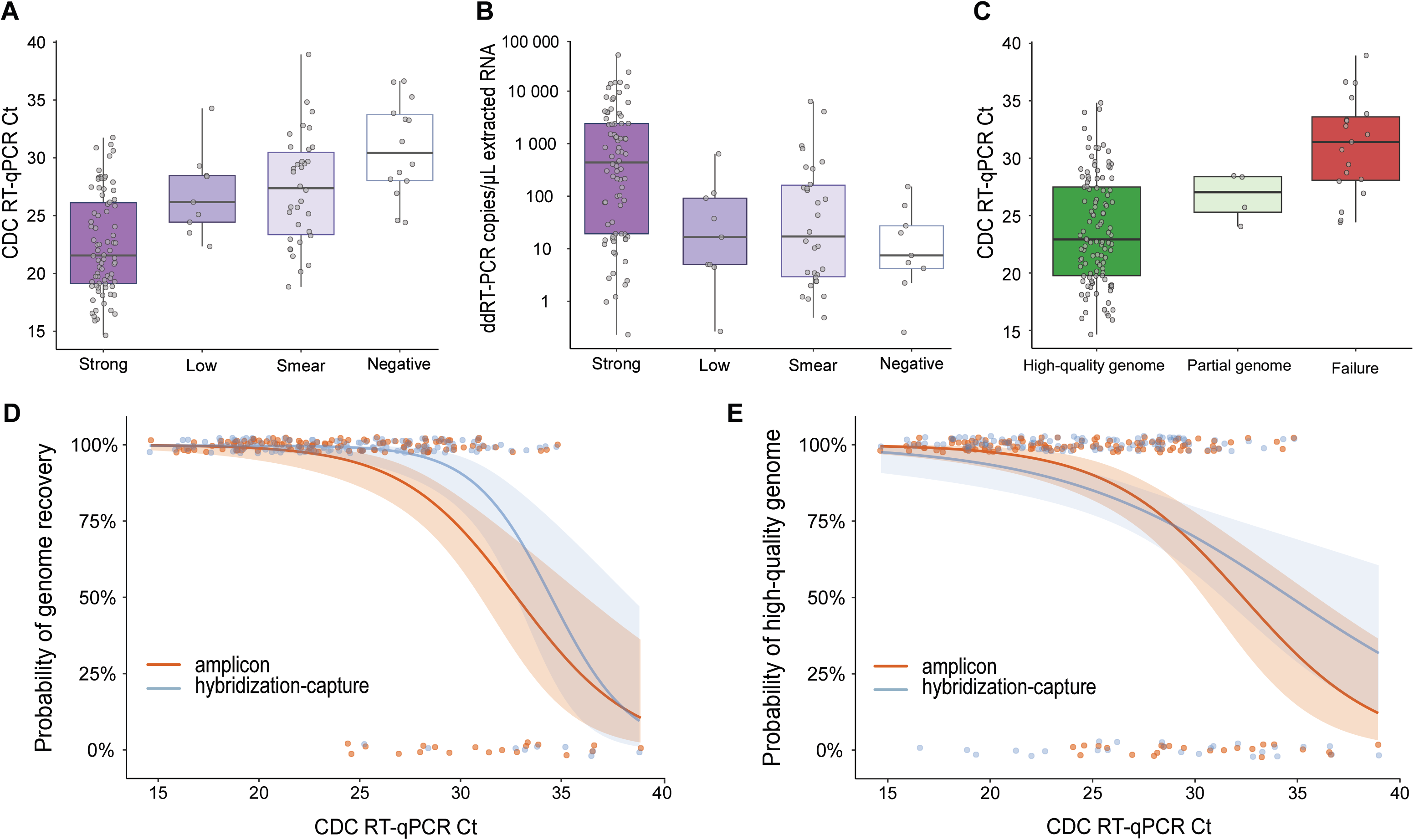
Association of viral burden and amplicon-band intensity with rhinovirus genome recovery. (A) CDC RT-qPCR Ct values and (B) ddRT-PCR viral RNA concentrations in extracted RNA, stratified by amplicon-band category. Points represent individual samples; boxes indicate the median and interquartile range, and whiskers extend to 1.5 times the interquartile range. The ddRT-PCR axis is logarithmically scaled. (C) CDC RT-qPCR Ct values according to single-amplicon sequencing outcome. (D) Logistic-regression estimates of the probability of recovering any genome, defined as a high-quality or partial genome rather than sequencing failure, as a function of CDC Ct value for single-amplicon and hybridization capture sequencing. (E) Logistic regression estimates for recovery of a high-quality genome versus a partial genome or failure. In D and E, points show observed binary outcomes with vertical jitter, lines show model-predicted probabilities, and shaded regions indicate 95% confidence intervals.

Genome recovery increased with viral abundance (Figure 2C). High-quality genomes were recovered from 95.7% (66/69) of specimens below a CDC Ct of 25 and from 30.0% (3/10) at Ct ≥33 (Figures 2 D/E). Predicted recovery by single-amplicon sequencing reached 90% at a CDC Ct of 25.1 (95% CI 22.7-27.4) for a high-quality genome and 26.5 (24.4-28.6) for any genome. Models fitted independently on the diagnostic platform scale placed similar thresholds at Panther Fusion Ct of 27.0 (24.3-29.8) and 27.9 (25.1-30.7). CDC Ct values predicted recovery of any genome at least as well for hybridization capture as for single-amplicon sequencing (AUC 0.92 versus 0.86) but predicted high-quality genome recovery less well (0.72 versus 0.84), because hybridization capture recovered partial sequence from low-abundance specimens (Supplementary Table S5).

Among specimens with a quantifiable ddRT-PCR result (126 out of 140), high-quality genomes were recovered from 34/34 specimens above 1,000 copies/µL extracted RNA, 30/31 between 100 and 1,000, 24/30 between 10 and 100, and 22/31 below 10 copies/µL. Absolute quantification was nonetheless the least precise predictor. Ten of the 22 single-amplicon failures had no quantifiable ddRT-PCR result and were therefore excluded, which truncated the low end of the measured range. The 90% threshold for a high-quality genome was 71 copies/µL of extracted RNA with a confidence interval that spanned more than an order of magnitude (95% CI 12 – 442). Estimates for every outcome, predictor and method are given in Supplementary Table S5.

### Single-amplicon sequencing recovered genetically diverse rhinovirus genomes

The single-amplicon approach recovered RV-A, RV-B, and RV-C genomes across 57 genotypes including 25 RV-A genotypes, 10 RV-B, and 22 RV-C (Supplementary Table S3). Single-infection was detected in 113 specimens and co-infections between two RV species or genotypes were detected in six specimens (RV-A68/RV-C7, RV-B69/RV-C12, RV-B104/RV-C23, RV-B104/RV-C36, and RV-C8/RV-C36). In addition, two specimens with non-RV enterovirus genomes were found: one single-infection (coxsackievirus A6) and one co-infection (RV-C31/coxsackievirus A16) (Supplementary Table S3). Cross-contamination did not explain co-infections: no pair of same-genotype genomes from different specimens in a sequencing run exceeded the 99.9% identity threshold used to flag possible carryover. Two of the RV genotypes involved, RV-C12 and RV-C23, and the coxsackievirus A16 were each recovered from a single specimen, leaving no potential source within the run. Nine patients contributed serial clinical specimens collected 8 to 69 days apart (Supplementary Table S3). Consensus genomes were identical between serial specimens from the same patient for every genotype recovered from both specimens, providing an internal confirmation of sample and sequence assignments (28).

From 116 specimens, paired amplicon and hybridization capture genomes were obtained and five corresponded to RV co-infections, yielding 121 matched specimen-genotype. Genotype assignments made by VP1 nucleotide identity were confirmed by maximum-likelihood phylogenetic analysis of VP1 for every genome and agreed between the two sequencing methods. The genotype RV-A80 (specimen 034157) was recovered by hybridization capture only. No recombinant genomes were detected in the sequenced cohort after comparison of phylogenetic associations inferred from VP1 and 3D genetic regions (phylogenetic analyses available online https://github.com/greninger-lab/RV_NGS_methods/).

### Consensus sequences were concordant between sequencing methods and reproducible within each method

Among 121 paired RV consensus genomes, eleven (9.1%) had at least one nucleotide substitution between methods, accounting for 25 canonical substitutions across the genome outside the 3′UTR-poly(A) boundary (Supplementary Table S6). Of these, 22 had a sequencing depth of ≥20x in both sequencing methods and could therefore be evaluated for intrahost diversity looking for the presence of the discrepancy as a minor allele in the other method. A total of 21/22 substitutions showed evidence of intrahost variation, defined as minor allele frequency ≥5% in at least one method (Figure 3). This revealed that only one sequence across 121 sequenced genomes had a discrepancy between methods that could not by intrahost diversity: specimen 175085 (RV-A18) (coordinate amplicon: 1114C / coordinate capture: 1466T). Additionally, 3′UTR-polyA boundary showed variability at 38 positions across 23 genomes, likely originated by polyT reverse primer mispriming, homopolymer slippage or read-mapping artifacts due to high-ambiguity in a region with a genetic context ‘TATAAAA’-like (Supplementary Table S6).

**Figure 3.**
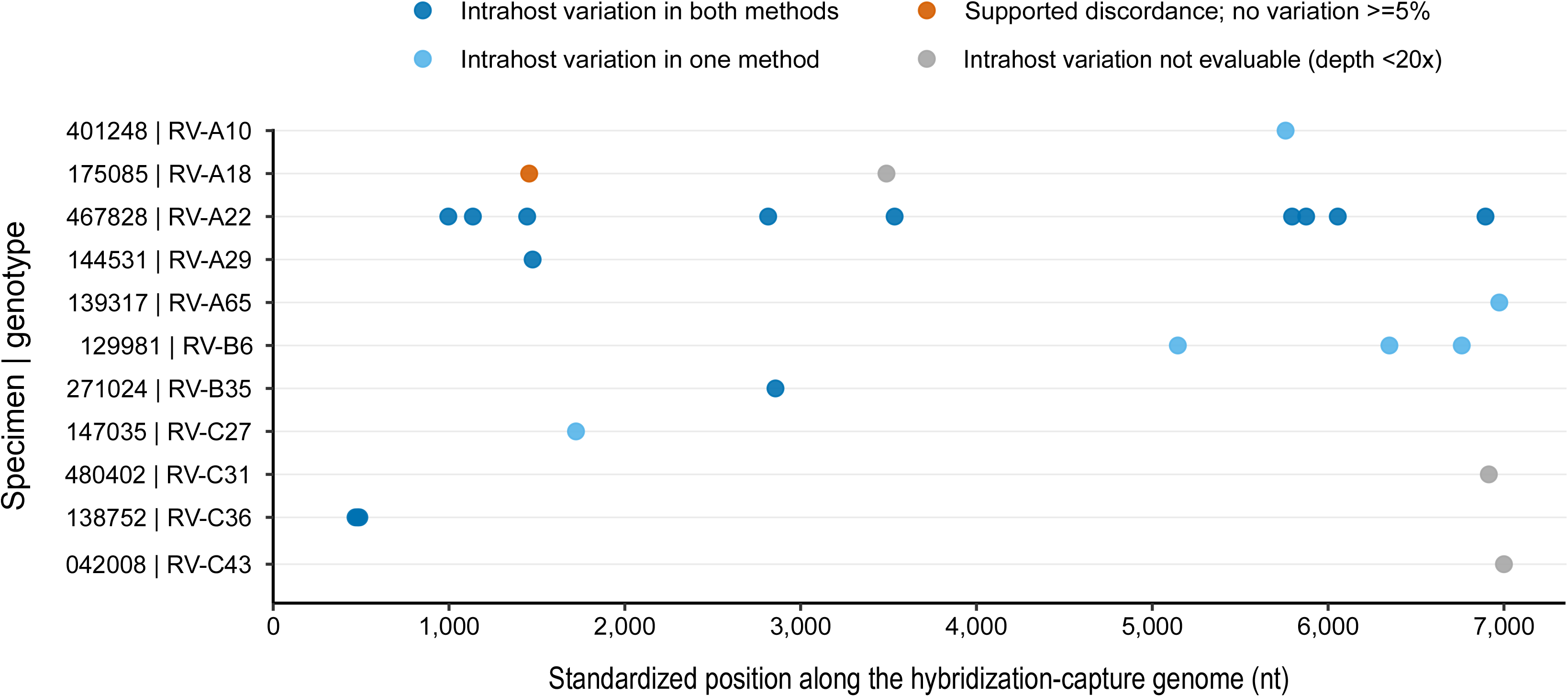
Genomic distribution and read-level support of canonical consensus differences between single-amplicon and hybridization capture sequencing. Each row represents a sample-genotype pair containing at least one canonical nucleotide difference between the single-amplicon sequencing and hybridization capture sequencing consensus genomes, and each point represents one canonical substitution. Positions were derived from the hybridization capture consensus since it is longer than amplicon-based consensus genome and standardized to a 7,000-nt genome to accommodate differences in genome length among genotypes. Colors indicate whether the two alternative consensus bases were supported as intrahost variants with a minor-allele frequency ≥5% in both methods, in one method only, or in neither method despite sufficient depth. Positions were considered evaluable when depth was above 20x in both methods; positions failing this criterion are shown with a grey dot.

Near-complete agreement between the two methods indicates that neither workflow systematically affected the consensus sequence, so we assessed whether a single run with either workflow is sufficient. We repeated the entire process (cDNA synthesis, PCR, library preparation, and sequencing) for 37 specimens by single-amplicon sequencing (19 with two replicates, 13 with three, and 5 with four) for which the complete amplicon was recovered in every case. Two of the tested specimens carried co-infections (416292, RV-B69/RV-C12; 352124, RV-A68/RV-C7), so these represented 39 sample-genotype groups and a total of 102 sequenced genomes. Median nucleotide identity among replicate consensus genomes was 99.99% (IQR 99.89-100%), and only two of the 39 sample-genotype groups contained substitutions in the consensus genomes, but the alternative allele was present with minor frequency (Supplementary Table S7). Twenty-three specimens were independently replicated by hybridization capture, with a median identity among replicates of 99.95% (IQR 99.54-99.98%) and three specimens contained consensus canonical discordances, also explained by intrahost variability with alleles present above 5% frequency (Supplementary Table S7). In both methods substitutions were confined to single specimens – 8/9 nucleotide discordances in specimen 467828 for single-amplicon and 7/9 in specimen 205980 for hybridization capture – rather than distributed across the cohort. Repeat execution of either workflow therefore returns essentially the same consensus sequence, and a single attempt is sufficient for routine use. Across the 16 sample-genotype groups replicated by both methods, median group-level pairwise concordance was similar for within-amplicon, within-capture and between-method comparisons (99.99%, 99.97% and 99.89%), and a Friedman test did not detect a difference (p-value = 0.36; all pairwise Wilcoxon test Benjamini-Hochberg-adjusted p ≥ 0.48).

### Analysis of mismatches in the f448 specific primer for single-amplicon sequencing

Hybridization capture sequencing extended through the first approximately 450 nt of the 5′UTR, which were not covered in the single-amplicon method (Figure 1A). This region includes the f448 forward-primer binding site, allowing direct examination of primer-template mismatches. Of 130 capture-derived consensus genomes that included sequence in the f448 binding site, four contained a single confirmed mismatch. Specimen 206743 (RV-A105) differed at primer position 23, where the primer contains T and the viral genome contains C (23:T>C). Specimens 257284, 400817, and 417427, all assigned to RV-A59, shared a single same mismatch (10:C>T). Single-amplicon sequencing nevertheless recovered specimens 206743 and 417427, which indicates that a single mismatch at primer position 10 or 23 may not necessarily preclude amplification.

Incompatibility with the f448 primer did not explain most single-amplicon sequencing failures, as 9 of the 11 failed specimens with an evaluable primer-binding site by hybridization capture sequencing had a conserved f448 sequence. Although failure was more frequent among specimens with a confirmed mismatch (2/4) than among those with a conserved site (9/126), the small number of mismatched specimens precluded a precise estimate of this association. The two primer-mismatched specimens that failed, 257284 and 400817, had no visible amplicon and relatively low ddRT-PCR-measured viral RNA concentrations (5.43 and 2.23 copies/µL extracted RNA, respectively), suggesting that factors other than primer-template compatibility, potentially including template abundance and RNA integrity, contributed to amplification failure.

### Single-amplicon sequencing reduced per-sample cost relative to hybridization capture and shotgun metagenomics

We compared per-sample costs for the single-amplicon and hybridization capture sequencing workflows from extracted RNA onward (Supplementary Table S8). For comparative purposes we also included the cost comparison for metagenomic sequencing with our previously published method (6). Per-sample reagent costs were $24.71 for single-amplicon sequencing, $32.70 for shotgun metagenomics, and $80.10 for hybridization capture. Tagmentation and indexing PCR accounted for $20.04 in both the single-amplicon and metagenomic workflows, so the pre-library preparation reagents that distinguishes them (reverse transcription, amplification, and cleanup) cost $4.67 and $12.66 per sample, respectively; in the capture workflow, library preparation is bundled into the enrichment kit and cannot be separated.

Single-amplicon and hybridization capture libraries were allocated 1 million reads per sample, whereas shotgun metagenomic libraries were allocated 2 million reads (which may not guarantee genome coverage, which cannot be verified until sequencing analysis is complete). At a rate of $5.00 per million reads, total per-sample costs were $29.71 for single-amplicon sequencing, $42.70 for metagenomics, and $85.10 for hybridization capture, a 2.9-fold reduction for single-amplicon sequencing relative to capture and a 1.4-fold reduction relative to metagenomics. The single-amplicon workflow also provides a pre-sequencing checkpoint that neither alternative offer: a visible amplicon band, confirmed by library quantification, identifies the specimens that will yield a genome (Supplementary Table S3), so specimens unlikely to yield a genome can be excluded before sequencing is performed. Personnel time differs as well, with single-amplicon and metagenomic library preparation each completed in one working day against three for hybridization capture, however labor costs were not considered in our calculations.

## Discussion

Here, we developed a single-round, single-amplicon RT-PCR and sequencing workflow that targets the enterovirus and rhinovirus polyprotein-coding region. The method recovered consensus sequences from 84% of specimens, covering 57 RV genotypes across rhinovirus A, B, and C. The same primer configuration recovered the polyprotein-coding sequence from every member of a 41-member enterovirus A–D panel. The protocol is simple and comparatively inexpensive to execute from a genomic surveillance perspective, especially when Ct values are available from the diagnostic platform involved in virus detection.

Genome-wide characterization of enteroviruses and rhinoviruses provides information that routine diagnostic testing and short genotyping targets do not. Complete coding-region sequences allow lineage assignment, placement within regional and global phylogenies, virus-like particle construction, and detection of recombination outside VP1. These capabilities have informed the emergence and spread of EV-D68, EV-A71, and poliovirus, and the lineages associated with outbreaks and severe disease (3–5, 29, 30). Rhinovirus genomic surveillance has likewise revealed extensive genotype co-circulation and recombinant lineages that detection alone does not resolve (6, 31, 32), although recombination frequency and genomic distribution differ among species and lineages (33). Because both primers bind conserved terminal sites, our amplification protocol is not expected to depend on the recombinant structure of the coding region.

Viral abundance was an important determinant of genome recovery for both single-amplicon and hybridization capture sequencing. RT-qPCR quantitation of enteroviruses and rhinoviruses can be variable (34), so we used two RT-qPCR assays, one on an FDA-cleared sample-to-answer diagnostic platform, in parallel with ddRT-PCR quantitation. The offset between the two RT-qPCR assays was consistent across RV species, arguing against genotype-specific probe mismatch and favoring a systematic difference in assay calibration or extraction efficiency. RNA fragmentation cannot easily be determined in advance, but the amplicon electrophoresis band in agarose gel and library quantification give an empirical readout of amplification success before sequencing.

Model-derived Ct and copy number values describe this cohort but should not be treated as universal acceptance thresholds. They depend on specimen processing, extraction volume, RT-qPCR assay, template integrity, and the definition of sequencing success, and they do not transfer to enterovirus-positive specimens quantified with an enterovirus-specific assay. These thresholds require local validation and prospective evaluation before confidently guiding routine specimen selection. Ct and copy number are upstream and predictive, allowing specimens to be prioritized before reagents are committed, whereas amplicon visualization with library quantification is downstream and, in this cohort, deterministic. Laboratories can therefore use their own Ct values to select specimens (predicted recovery of a high-quality genome reached 90% at a CDC Ct of 25.1 or a Panther Fusion Ct of 27.0 in this cohort), and the gel and library quantification as the decision point before a sequencing slot is committed. Recovery declined to 30.0% at a CDC Ct ≥33, where the gel was the more reliable guide. Specimens without a visible amplicon remained negative when re-tested once from the original extract; where lineage assignment is required, conventional VP1 genotyping can be attempted. Hybridization capture recovered sequence from more specimens but returned three times as many partial genomes (19 versus 6), so single-amplicon sequencing is preferable when coding-complete genomes are the objective.

A recent expert view from the European non-polio enterovirus network identifies the length-sensitivity trade-off as the central constraint of targeted enterovirus sequencing, because longer amplicons are more informative but reduce PCR sensitivity (12). Two prior protocols have described recovery of the enterovirus coding region from a single amplicon, but these have never been evaluated for rhinovirus genome recovery (18, 19). The inner forward primer of the nested RT-PCR described by Isaacs et al. anneals at the same conserved 5′ UTR position as f448, but is 5 nt shorter, and differs at two positions where f448 incorporates ambiguity bases (9:Y>C and 19:Y>C). Chien et al. paired a 5′ UTR forward primer overlapping f448 across 16 nt with a tag introduced during reverse transcription (18, 19). These primers were derived independently (ours from the CDC rhinovirus RT-qPCR probe, theirs from earlier enterovirus primer sets), and their convergence on the same position within the IRES supports this locus as a priming site for genus-wide amplification. Although the CDC RT-qPCR probe was design as RV-specific, the site that f448 targets is conserved across the genus: the primer amplified coxsackieviruses A6 and A16 and the enterovirus A-D panel. Of note, our IRES primer ended f448 at the last position conserved across the genus. Neither protocol was characterized on rhinoviruses: Isaacs et al reports a single RV-A56 amplification as data not shown, and Chien et al. evaluated only on cultured isolates. Here, the method was evaluated in cultured isolates but also in primary diagnostic specimens spanning the full viral load range and benchmarked against hybridization capture in the same specimens.

Across the six nucleotides by which the primer of Chien et al. extends further 3′, the proportion of human enterovirus and rhinovirus NCBI sequences (n = 20,972) carrying a substitution or indel was strongly virus-dependent, ranging from 0.6% in coxsackievirus A10 and 1.9% in coxsackievirus A16 to 78.7% in coxsackievirus A24. It was highest in rhinoviruses, at 85.8% of RV-A, 99.3% of RV-B and 63.8% of RV-C sequences, or 79.4% of rhinovirus sequences overall against 28.2% of enterovirus sequences (analysis available at https://github.com/greninger-lab/RV_NGS_methods). Whether mispairing in this window reduces amplification cannot be resolved from sequence alone, and Chien et al. amplified types in which such mismatches are common; for an assay intended to cover the genus, the relevant property is the worst case rather than the average match.

The nested format proposed by Isaacs et al. requires two sequential reactions and four virus-specific priming sites, two in the 5′UTR and two in the 3′UTR. In contrast, our single-round of amplification depends on one virus-specific site and reads through to the poly(A) junction, recovering the complete 3′ UTR. The Isaacs et al. inner forward primer, equivalent to f448, matched exactly 82.8% (17,357/20,972) of the analyzed sequences, while 17.2% (3,605/20,972) carried one to three substitutions. Most of these differences (3,480 sequences) involved 19:Y>C, where the alternative nucleotide is present in 71.8% of RV-A and 31.1% of RV-C sequences, compared with 3.3% of EV-C and <0.5% in the remaining species. Although nested PCR can improve analytical sensitivity, successful amplification depends on the generation of suitable products during the first reaction. The outer forward primer required for the first round of Isaacs et al. protocol matched exactly 41.5% (6,016/14,513) of sequences with an evaluable binding site, while 58.4% (8,481/14,513) carried one to three substitutions. Exact match frequencies varied by species from 9.5% EV-B (171/1,806) to 78.8% RV-A (1463/1,857). These analyses only examine primer site conservation rather than amplification performance, which also depends on mismatch position and type, polymerase properties, cycling conditions, and template abundance.

The enterovirus panel provides initial evidence that our primer configuration is broadly reactive beyond rhinovirus A-C, consistent with earlier work showing that near-full-length or coding-complete enterovirus sequences can be generated by targeted amplification (15, 18, 19). In the clinical cohort, genomes were recovered from two of the four specimens carrying a confirmed f448 mismatch, including one at primer position 23, within the three terminal nucleotides where mispairing is most likely to impair extension, and both specimens that failed also had low measured viral load. Whether the 22:C>A substitution that predominates in EV-C104, EV-C105 and EV-C109 is similarly tolerated remains untested, because these three types were not available to us.

The amplicon starts at f448 and omits the first approximately 450 nt of the 5′ UTR. The single-round design gains simplicity and highly uniform depth, at the cost of a product that is not a complete genome, cannot be used for reverse genetics cloning and cannot reveal 5′ UTR recombination breakpoints, which matters for enterovirus C and poliovirus surveillance.

This study has limitations. It was retrospective, single center and enriched for rhinovirus-positive respiratory specimens. No experimental attempt was made in enterovirus-negative or unrelated clinical specimens, and it is unclear whether the approach would amplify enterovirus genomes from stool or wastewater. The enterovirus panel consisted of cultured isolates, and the two enterovirus-positive clinical specimens were incidental detections in a cohort selected as RV-positive, so performance in primary enterovirus specimens at low abundance, with degraded RNA or at divergent primer-binding sites remains to be shown. A single product of approximately 7 kb is well suited to long-read sequencing (19), which we did not attempt; any such implementation would need a pipeline able to detect same-species co-infections and flag potential cross-contamination, as ours does. In addition, the amplicon recovered omits the first approximately 450 nt of the enterovirus/rhinovirus genome, which means it cannot be for reverse genetics cloning and cannot reveal 5′ UTR recombination breakpoints, which matters for enterovirus C and poliovirus surveillance.

In summary, single-amplicon sequencing recovered reproducible polyprotein-coding-region consensus sequences across a diverse rhinovirus cohort and enterovirus panel, with a simple amplification scheme, highly uniform depth and an informative pre-library quality control check. Given the impressively high burden of enteroviruses and rhinoviruses, single-amplicon sequencing could provide a practical approach for large-scale lineage characterization, genomic surveillance, and investigation of transmission and recombination.

## Acknowledgements

We thank Dr. Gabriel Parra for valuable insights in single-amplicon sequencing methods and Dr. Jonathan Reed for helpful comments on the manuscript. Claude Opus 5.5 (Anthropic) was used in October 2026 to edit the grammar, clarity, and style of text written by the authors. No manuscript content was generated by the AI tools. ALG reports contract testing to UW from Abbott, Cepheid, Novavax, Pfizer, Janssen, Assembly Biosciences, Aicuris, Innovative Molecules, and Hologic, research support from Gilead, personal fees from Arisan Therapeutics, outside of the described work. This work was supported by NIAID R21 AI185726 and the University of Washington Department of Laboratory Medicine and Pathology. The remaining authors report no conflicts of interest. The funders had no role in study design, data collection and interpretation, or the decision to submit the work for publication.

